# polyRAD: Genotype calling with uncertainty from sequencing data in polyploids and diploids

**DOI:** 10.1101/380899

**Authors:** Lindsay V. Clark, Alexander E. Lipka, Erik J. Sacks

## Abstract

Low or uneven read depth is a common limitation of genotyping-by-sequencing (GBS) and restriction site-associated DNA sequencing (RAD-seq), resulting in high missing data rates, heterozygotes miscalled as homozygotes, and uncertainty of allele copy number in heterozygous polyploids. Bayesian genotype calling can mitigate these issues, but previously has only been implemented in software that requires a reference genome or uses priors that may be inappropriate for the population. Here we present several novel Bayesian algorithms that estimate genotype posterior probabilities, all of which are implemented in a new R package, polyRAD. Appropriate priors can be specified for mapping populations, populations in Hardy-Weinberg equilibrium, or structured populations, and in each case can be informed by genotypes at linked markers. The polyRAD software imports read depth from several existing pipelines, and outputs continuous or discrete numerical genotypes suitable for analyses such as genome-wide association and genomic prediction.

## Introduction

Approximately 70% of vascular plant species are recent polyploids, yet genomic resources and bioinformatics tools for polyploids typically lag behind those for diploids (Moghe and Shiu 2014; Renny-Byfield and Wendel 2014; Bourke *et al*. 2018b). Reduced representation DNA sequencing methods, such as genotyping-by-sequencing (GBS) and restriction site-associated DNA sequencing (RAD-seq), have made high-density genotyping considerably more accessible and affordable (Poland and Rife 2012; Davey *et al*. 2013). However, the two most popular pipelines for processing GBS and RAD-seq data, Stacks (Catchen *et al*. 2013) and TASSEL (Glaubitz *et al*. 2014), do not output polyploid genotypes. Though pipelines for polyploids are available, each have limitations that prevent their general application. For example, the UNEAK pipeline is designed for diploidized polyploids only (Lu *et al*. 2013). HaploTag is specialized for self-fertilizing polyploids (Tinker *et al*. 2016). FreeBayes and GATK can output polyploid genotypes, but require a reference genome (McKenna *et al*. 2010; Garrison and Marth 2012). The software EBG imports read depth from other pipelines to estimate auto- or allopolyploid genotypes (Blischak *et al*. 2018) but requires allele frequency estimations from the parent species for allopolyploids. The R package updog estimates polyploid genotypes from read depth, modeling preferential pairing and accounting for multiple technical issues that can arise with sequencing data, and can output posterior mean genotypes reflecting genotype uncertainty (Gerard *et al*. 2018), but requires excessive amounts of computational time to run. SuperMASSA (Serang *et al*. 2012) and fitPoly (Voorrips *et al*. 2011) were originally designed for calling polyploid genotypes from fluorescence-based SNP assays and have been adapted for sequencing data, but fail to call genotypes when low read depth results in high variance of read depth ratios. Thus, important staple crops such as wheat, potato, sweet potato, yam, and plantain are underserved by existing genotyping software, limiting our ability to perform marker-assisted selection, while yield increases from breeding are not keeping pace with projected food demands (Ray *et al*. 2013).

We present a new R package, polyRAD, for genotype estimation from read depth in polyploids and diploids. The software polyRAD is designed on the principle originally proposed by Li (2011) that it is not necessary to call genotypes with complete certainty in order to make useful inferences from sequencing data. Initially, SNP discovery is performed by other software such as TASSEL (Glaubitz *et al*. 2014) or Stacks (Catchen *et al*. 2013), with or without a reference genome, then allelic read depth is imported into polyRAD from those pipelines or the read counting software TagDigger (Clark and Sacks 2016). In polyRAD, one or several ploidies can be specified, including any level of auto- and/or allopolyploidy, allowing inheritance modes to vary across the genome. Genotype probabilities are estimated by polyRAD under a Bayesian framework, where priors are based on mapping population design, Hardy-Weinberg equilibrium (HWE), or population structure, with or without linkage disequilibrium (LD) and/or self-fertilization. Multi-allelic loci (haplotypes) are allowed, and are in fact encouraged because LD within the span of one RAD tag is not informative for genotype imputation. In addition to exporting the most probable genotype for each individual and locus, continuous numerical genotypes can be exported reflecting the relative probabilities of all possible allele copy numbers, and can then be used for genome-wide association or genomic prediction in software such as GAPIT (Lipka *et al*. 2012), FarmCPU (Liu *et al*. 2016b), TASSEL (Bradbury *et al*. 2007), or rrBLUP (Endelman 2011). Discrete genotypes can also be exported for polymapR (Bourke *et al*. 2018a). polyRAD is the first Bayesian genotype caller to incorporate population structure and multiple inheritance modes, as well as the first with an option for mapping population designs other than F1 and F2. It is available at https://github.com/lvclark/polyRAD and https://CRAN.R-project.org/package=polyRAD.

## Methods

### Overview

polyRAD implements Bayesian genotype estimation, similar to that proposed and implemented by several other groups (Li 2011; Nielsen *et al*. 2011; Garrison and Marth 2012; Korneliussen *et al*. 2014; Maruki and Lynch 2017; Gerard *et al*. 2018; Blischak *et al*. 2018). In all polyRAD pipelines, genotype prior probabilities (*P*(*G_i_*)) represent, for a given allele and individual, the probability that *i* is the true allele copy number, before taking allelic read depth into account. Genotype prior probabilities are specified from population parameters, and optionally from genotypes at linked markers (see Supplementary Methods).

For a given individual and locus, consider every sequencing read to be a Bernoulli trial, where the read either matches a given allele (success) or some other allele (failure). The probability of success is:

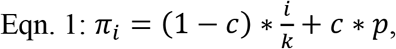

where *c* is the cross-contamination rate, *i* is the allele copy number in the genotype, *k* is the ploidy, and *p* is the allele frequency in the population. For GBS and RAD-seq data, *c* is estimated by including a negative control in library preparation, i.e. of the set of ligation reactions with barcoded adapters, one that has no genomic DNA added. The sequence read depth for this blank barcode is then divided by the mean read depth of non-blank barcodes in order to estimate *c*. In practice we have found *c* to typically be 1/1000 (unpublished data), and expect it to be more substantial than errors arising from the sequencing technology, which will tend to produce haplotypes not found elsewhere in the data set. The *c* parameter is important for identifying homozygotes that could otherwise be misidentified as heterozygotes.

Gerard et al. (2018) observed overdispersion in the distribution of sequence read depth, indicating that in reality *π_i_* varies from sample to sample. We have observed the same in our datasets, likely due to factors such as differing contamination rates among samples, restriction cut site variation, and differences in size selection among libraries. Therefore, following Gerard et al. (2018), we model allelic read depth as following a beta-binomial distribution rather than a binomial distribution. For every possible allele copy number at a given locus and individual, the following equation is used to estimate the likelihood of the observed read depth using the beta-binomial probability mass function:

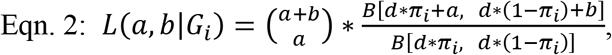

where *a* is the number of reads for a given allele at a given locus, *b* is the number of reads for other alleles at that locus, *G_i_* is the state in which a locus has *i* copies of a given allele, *B* is the beta function, and *d* is the overdispersion parameter. The parameter *d* is set to nine by default given our observations of overdispersion in empirical data, and can be increased to model less overdispersion and vice versa. The function *TestOverdispersion* is included in polyRAD to assist the user in determining the optimal value of *d*.

From the priors and likelihoods, a posterior probability can then be estimated for each possible allele copy number for each individual and allele using Bayes’ theorem (Shiryaev 2011):

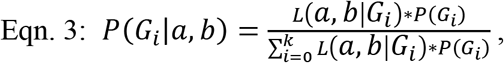

where all terms are as previously described.

Bayesian genotype estimation allows correction of genotyping errors in diploids and polyploids, i.e. when an individual is truly heterozygous but only one allele was sequenced, or when an individual appears heterozygous due to sequencing error or contamination but is truly homozygous. It also enables estimation of allele dosage in heterozygous polyploid genotypes. Moreover, genotype posterior probabilities are more influenced by priors when read depth is low, and by genotype likelihoods derived from allelic read depth when read depth is high. When read depth is zero for a given individual and locus, genotype posterior probabilities are equal to priors, and thus missing and non-missing data are handled within one coherent paradigm. It is therefore not necessary to impute missing genotypes in a second step if the priors are sufficiently informative.

For export to other software, as well as iteration within the polyRAD pipelines, a given allele’s posterior mean genotype (*pmg*) is a mean of the number of copies of that allele, with the posterior genotype probabilities (Eqn. 3) serving as weights, as in Guan and Stephens (2008). Thus, for an individual and allele, *pmg* is calculated as:

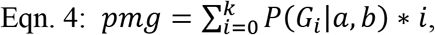

where all terms are as previously described. Additional details and equations for specification of prior genotype probabilities and estimation of other parameters are provided in Supplementary Materials. A flow chart of how this Bayesian genotypic estimation is implemented into polyRAD is displayed in Fig. 1. In brief, for mapping populations, genotype priors are specified based on parental genotypes and progeny allele frequencies, and all parameters are estimated once. For diversity panels, genotype priors are adjusted and parameters re-estimated iteratively until allele frequencies converge. Source code is available at https://github.com/lvclark/polyRAD, archived at Zenodo (doi: 10.5281/zenodo.1143744).

**Fig. 1.**
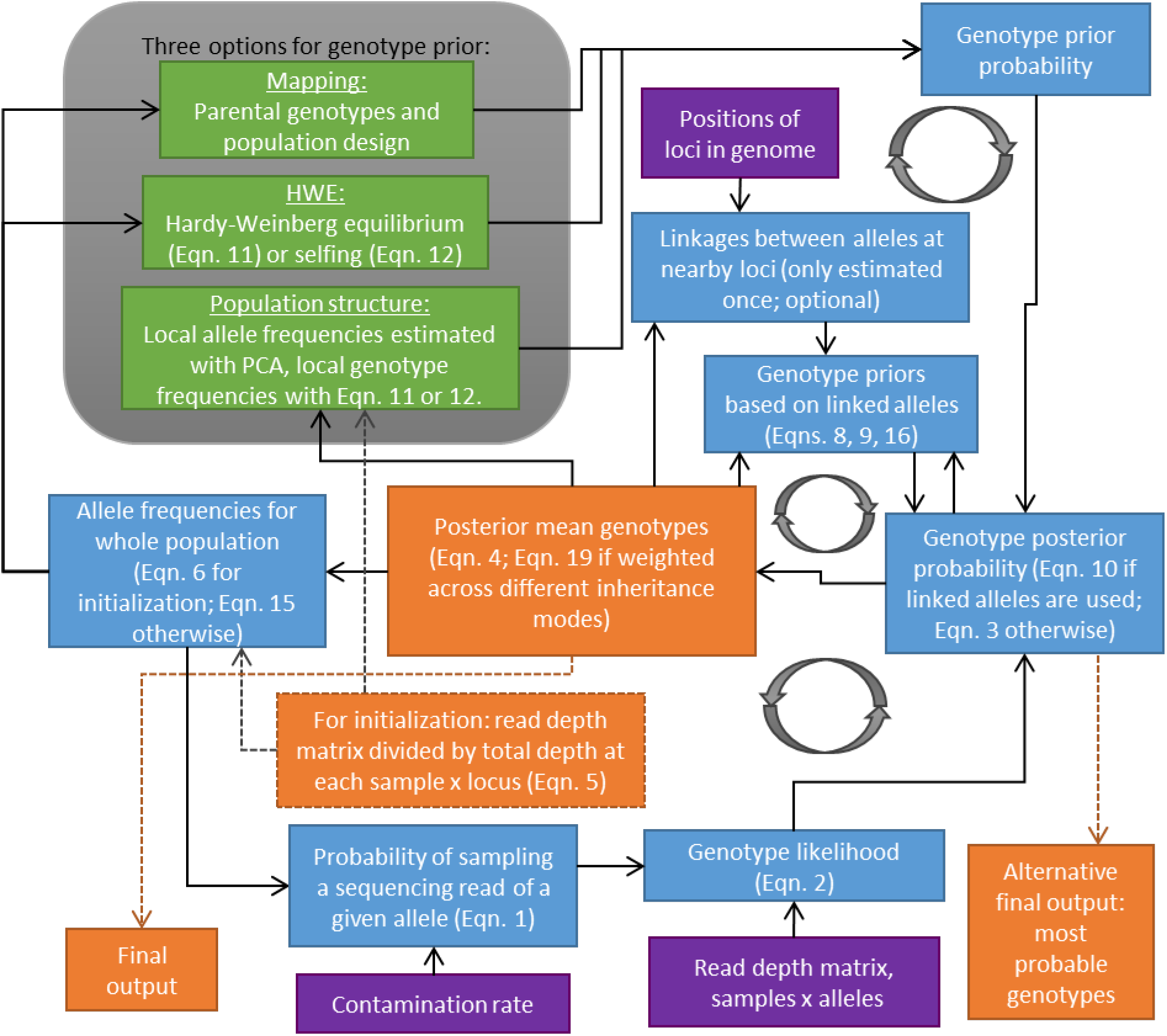
Overview of polyRAD algorithms for genotype estimation. Genotype posterior probabilities are estimated iteratively until allele frequencies converge, except in the case of mapping populations, where allele frequencies are only estimated once. Purple boxes indicate inputs to the pipeline (read depth, contamination rate, and optionally, genomic positions of loci). Blue boxes indicate estimated parameters (allele frequencies, genotype likelihoods and prior and posterior probabilities, linkage between alleles, and probability of sampling each allele). Green boxes indicate alternative methodologies for genotype prior probability estimation (mapping, HWE, and population structure). Priors for the HWE and population structure models can be adjusted for self-fertilization according to de Silva et al. (2005). Orange boxes indicate sample × allele matrices indicating approximate allele copy number. Dashed arrows indicate steps that happen only once at the beginning or end of the pipeline, whereas solid arrows indicate iterative steps. Circular arrows highlight cycles of iteration. Eqns. 1-4 are provided in the main manuscript, and Eqns. 5-19 are provided in Supplemental Materials.

### Example use

Executable examples are provided in the vignette and manual distributed with polyRAD. Here we provide an additional brief example. Box 1 illustrates the use of polyRAD on a diversity panel of a generic tetraploid species with a reference genome. Tools from the Bioconductor package VariantAnnotation (Obenchain *et al*. 2014) are used by the polyRAD function *VCF2RADdata* for import of a VCF file to the polyRAD-specific “RADdata” format. SNP filtering criteria are specified with the *min.ind.with.reads* and *min.ind.with.minor.allele* arguments to indicate the minimum number of individuals that must have more than zero reads of a locus, and the minimum number of individuals that must have reads of the minor allele, respectively. The *possiblePloidies* argument indicates that the inheritance mode could be allotetraploid (*c(2,2*)) or autotetraploid (*4*). Any ploidy may be specified with *possiblePloidies*, for example *8* for auto-octoploid, with the only limitation that all subgenomes in an allopolyploid must have the same ploidy. By default, *VCF2RADdata* groups SNP alleles into haplotypes that appear to have come from the same RAD tag, the size of which is specified by *tagsize*, in basepairs. Negative controls are indicated with *SetBlankTaxa*, and the contamination rate is estimated with *EstimateContaminationRate*. The function *IteratePopStructLD* is then used for genotype estimation, taking both population structure and LD into account. The probabilistic principal components analysis method from the Bioconductor package pcaMethods (Stacklies *et al*. 2007) is used internally by *IteratePopStructLD* in order to estimate population structure. The *LDdist* argument indicates the distance in basepairs within which to search for alleles at other loci that can help predict copy number of a given allele. Once genotype posterior probabilities are estimated, other parameters are cleared from memory using the *StripDown* function. Continuous numerical genotypes are then formatted for GAPIT (Lipka *et al*. 2012) using the *ExportGAPIT* function. Alternative functions are listed in Table 1. A very similar script could be used for a species without a reference genome, with *IteratePopStruct* in place of *IteratePopStructLD*, and a different import function for the appropriate non-reference pipeline.

#### Box 1. Example R script using polyRAD. Read depth is imported from a VCF file, genotypes are estimated using population structure and LD, and continuous numerical genotypes are formatted for GAPIT.

~~~
library(polyRAD)
library(VariantAnnotation)
# prepare the VCF file for import
myvcf <- “somegenotypes.vcf”
myvcfbg <- bgzip(myvcf)
indexTabix(myvcfbz, format = “vcf”)
# import VCF into a RADdata object
myRAD <- VCF2RADdata(myvcfbg,
                            tagsize = 64,
                            min.ind.with.reads = 300,
                            min.ind.with.minor.allele = 15,
                            possiblePloidies = list(c(2,2), 4))
# estimate contamination rate
myRAD <- SetBlankTaxa(myRAD, c(“blank1”, “blank2”))
myRAD <- EstimateContaminationRate(myRAD)
# genotype estimation with pop. structure pipeline
myRAD <- IteratePopStructLD(myRAD, LDdist = 5e4)
# free up memory
myRAD <- StripDown(myRAD)
# export for GAPIT
myGM_GD <- ExportGAPIT(myRAD)
~~~

**Table 1.** Overview of main polyRAD functions.

|  |  |
| --- | --- |
| **Import functions** |  |
| VCF2RADdata | Imports any VCF with an allelic read depth (AD) field, such as those exported by TASSEL-GBSv2 or GATK. |
| readTagDigger | Imports CSV file of read depth output by TagDigger. |
| readStacks | Reads catalog and matches files from Stacks. |
| readTASSELGBSv2 | Reads SAM and TagTaxaDist files from TASSEL-GBSv2. |
| readHMC | Reads files output by UNEAK. |
| **Genotype estimation functions** |  |
| PipelineMapping2Parents | For mapping populations with any number of generations of backcrossing, intermating, and/or selfing. |
| IterateHWE | For diversity panels without population structure.<sup>a</sup> |
| IterateHWE_LD | For diversity panels with LD and without population structure.<sup>a</sup> |
| IteratePopStruct | For diversity panels with population structure.<sup>a</sup> |
| IteratePopStructLD | For diversity panels with population structure and LD.<sup>a</sup> |
| **Export functions** |  |
| ExportGAPIT | Format genotypes for the *GD* and *GM* arguments of GAPIT or FarmCPU. |
| Export_rrBLUP_Amat | Format genotypes for the *A.mat* function in rrBLUP. |
| Export_rrBLUP_GWAS | Format genotypes for the *GWAS* function in rrBLUP. |
| Export_TASSEL_Numeric | Write file formatted for TASSEL with continuous numeric genotypes. |
| Export_polymapR | Format genotypes for the polymapR package. |
| GetWeightedMeanGenotypes | Create a matrix of continuous numeric genotypes. |
| GetProbableGenotypes | Create a matrix of discrete genotypes, indicating the most probable genotype for each individual and allele. |

<sup>a</sup>The rate of self-fertilization can be specified for self-compatible plant species.

### Testing

To test the accuracy of polyRAD, we used datasets from three previously studied populations: 1) RAD-seq data and GoldenGate SNP genotypes from a diversity panel (n = 565) of the outcrossing, diploidized allotetraploid grass *Miscanthus sinensis* (Clark *et al*. 2014), 2) RAD-seq data and GoldenGate SNP genotypes from a bi-parental F_1_ mapping population (n = 275) of *M. sinensis* (Liu *et al*. 2016a), and 3) SNP array genotypes from a biparental F1 mapping population of autotetraploid potato (n = 238) (da Silva *et al*. 2017). Allelic read depth at simulated RAD-seq markers was generated from the GoldenGate or SNP array genotypes, with overall locus depth drawn from a gamma distribution to resemble depth of actual RAD-seq markers (shape = 2 and scale = 5). The read depth for an individual genotype was also sampled from a gamma distribution, with the shape equal to the locus depth divided by 10, and scale = 10. The read depth for each allele was then sampled from the beta-binomial distribution as described in Eqn. 2, with *d* = 9 and *c* = 0.001. The *M. sinensis* diversity panel included 395 GoldenGate markers, plus real RAD-seq data for those same individuals across 3290 tag locations within 20 kb of any GoldenGate markers, called with the TASSEL GBS v2 pipeline (Glaubitz *et al*. 2014) using the *M. sinensis* v7.1 reference genome (DOE-JGI, http://phytozome.jgi.doe.gov/). Additionally, to test the effect of ploidy within the *M. sinensis* diversity panel, tetraploidy was simulated by summing GoldenGate genotypes and RAD-seq read depth of each individual with the individual with the most similar read depth to it out of the ten individuals most closely related to it. The *M. sinensis* mapping population included 241 GoldenGate markers genotyped across 83 individuals, plus 3062 RAD-seq markers called with the UNEAK pipeline (Lu *et al*. 2013) across those 83 individuals plus an additional 192 individuals. The potato mapping population included genotypes at 2538 markers. Additional simulations using data from diversity panels of soybean (Song *et al*. 2015), apple (Chagné *et al*. 2012), and potato (Hamilton *et al*. 2011) are presented in Figs. S1-S4. In each population, the simulated and real RAD-seq data was used for genotype calling with polyRAD, EBG (Blischak *et al*. 2018), updog (Gerard *et al*. 2018), and fitPoly (Voorrips *et al*. 2011), and missing genotypes from the EBG output were imputed with LinkImpute (Money *et al*. 2015) and/or rrBLUP (Endelman 2011) as appropriate. To estimate the accuracy of genotype calling and imputation, the root mean squared error (RMSE) was calculated between numeric genotypes (ranging from zero to the ploidy) at each simulated RAD-seq marker and at the GoldenGate or SNP array marker from which it was derived. Data and scripts for analysis are available at https://doi.org/10.13012/B2IDB-9729830_V2. Supplementary text, equations, and figures have been deposited at Figshare: https://doi.org/10.25387/g3.xxxxxxx.

## Results and discussion

### Accuracy of polyRAD

In the *M. sinensis* diversity panel, polyRAD showed improved genotype accuracy over the HWE, disequilibrium, and GATK methods implemented in EBG, as well as fitPoly, particularly at low read depths (Figs. 2A and 3A). polyRAD also showed a modest improvement in accuracy across all read depths as compared to updog (Figs. 2A and 3A) while needing approximately two orders of magnitude less processing time than updog. Under the HWE model in polyRAD with discrete genotypes output, errors in genotypes with more than zero reads were similar to those from the HWE model of EBG in both diploid and tetraploid systems (Figs. 2A and 3A). However, when priors in polyRAD were based on population structure, errors decreased, particularly in tetraploids and at low read depth (Figs. 2A and 3A). In diploids and tetraploids respectively using the polyRAD population structure model with discrete genotypes, error (RMSE) was reduced by 14.6% (SE 1.0%) and 23.5% (SE 0.6%) relative to the GATK model, by 10.5% (SE 0.9%) and 11.8% (SE 0.5%) relative to the EBG HWE model, by 26.0% (SE 1.2%) and 25.6% (SE 0.6%) relative fitPoly, and by 8.0% (SE 1.0%) and 18.0% (SE 0.7%) relative to discrete genotype output by the updog “norm” model. Given the known population structure in *M. sinensis* (Clark *et al*. 2014), it is unsurprising that a population structure-aware genotyping method would be more accurate than those based on HWE or otherwise not accounting for population structure. For genotypes with zero reads, imputation was most accurate when it accounted for population structure, using either polyRAD or rrBLUP (Fig. 2B and 3B). Although modeling LD did not improve accuracy in *M. sinensis* (Figs. 2 and 3), likely due to low LD as a result of outcrossing (Slavov *et al*. 2014), modeling LD did improve accuracy in wild soybean, apple, and a simulated inbreeding allohexaploid (Figs. S1, S2, and S3, and Supporting Results). In a diversity panel of tetraploid potato, accuracy was improved by modeling population structure but not LD (Fig. S4 and Supporting Results).

**Fig. 2.**
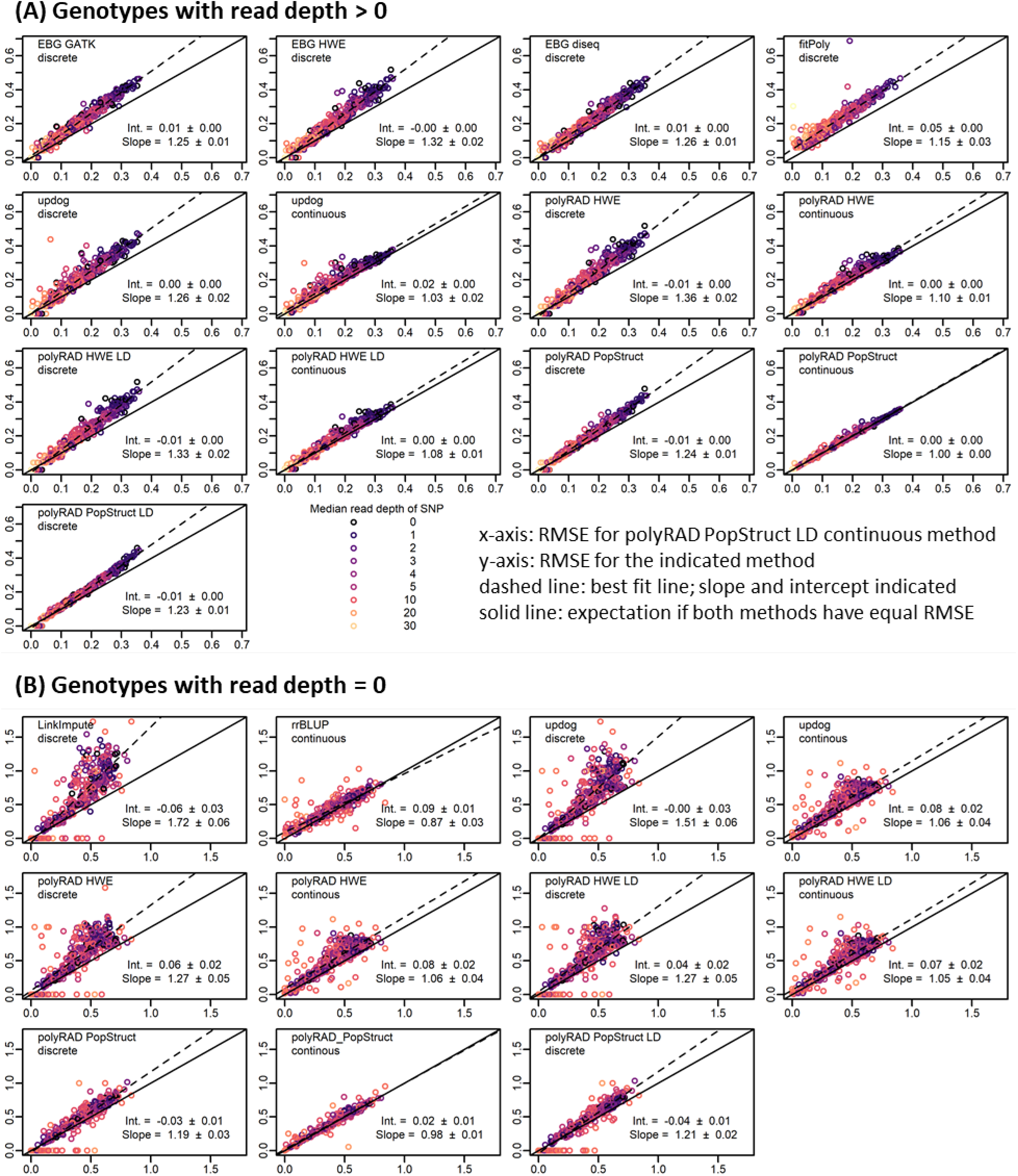
Genotyping error of EBG, fitPoly, updog, polyRAD, LinkImpute, and rrBLUP in a diversity panel of 565 diploid *Miscanthus sinensis*. The benefits of incorporating population structure into the genotyping model and using continuous rather than discrete genotypes are illustrated. Genotypes were coded on a scale of 0 to 2. Root mean squared error (RMSE) was calculated between actual genotypes and genotypes ascertained from simulated RAD-seq reads at 395 SNP markers (lower RMSE = higher accuracy). Each point represents one SNP. Median read depth is indicated by color, including genotypes with zero reads. The RMSE for continuous genotypes output by the polyRAD PopStruct LD method is shown on the x-axis, and the RMSE of other methods and types of genotypes (continuous or discrete) is shown on the y-axis. The dashed line indicates the ordinary least-squares regression with slope and intercept estimates, with standard errors. The “norm” model was used with updog. (A) RMSE calculated using only genotypes with more than zero reads. (B) RMSE calculated using only genotypes with zero reads, by genotyping or imputation method and genotype type.

**Fig. 3.**
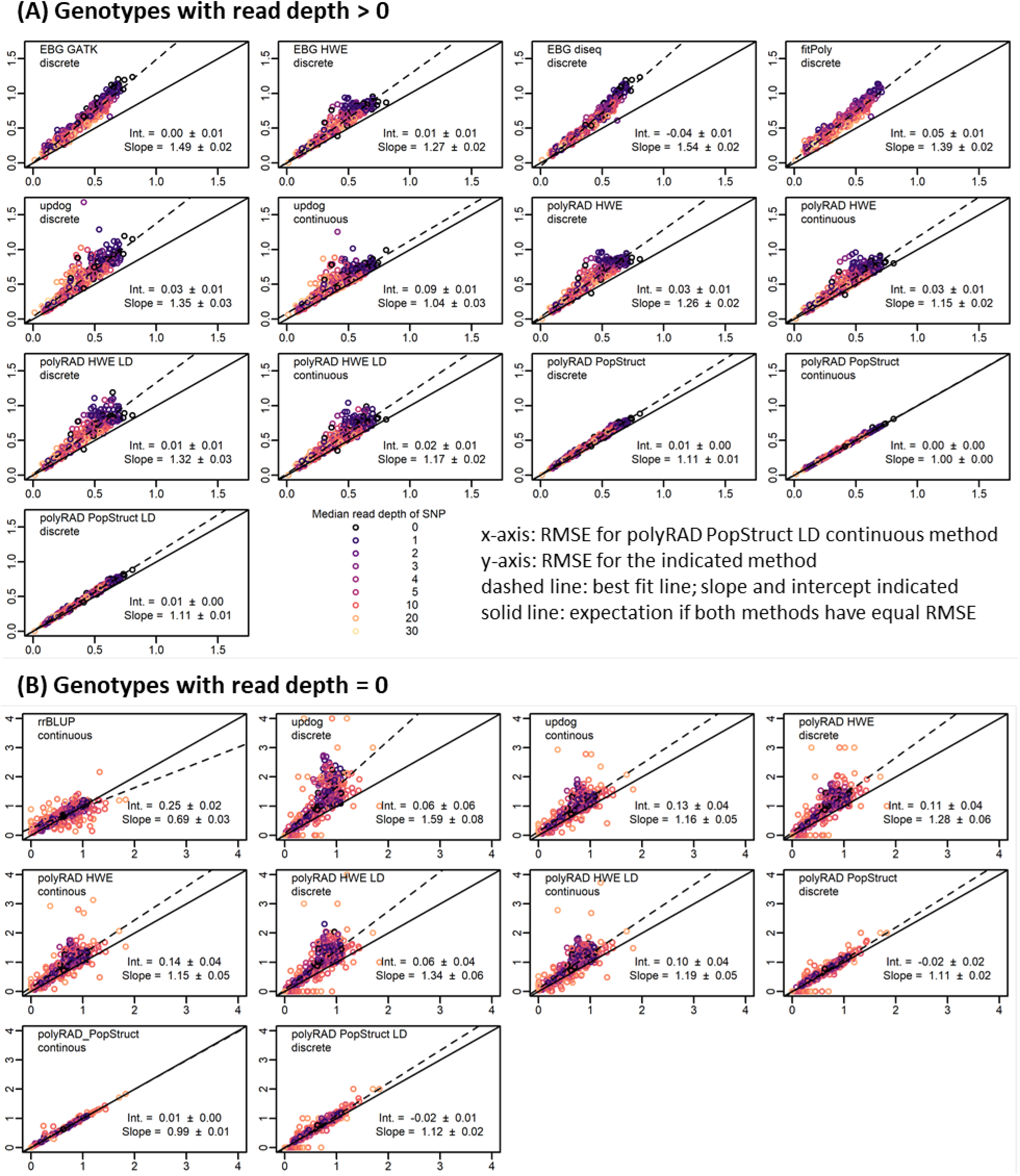
Genotyping error of EBG, fitPoly, updog, polyRAD, and rrBLUP in a simulated tetraploid diversity panel derived from genotypes of 565 diploid *Miscanthus sinensis*. The benefits of incorporating population structure into the genotyping model and using continuous rather than discrete genotypes are illustrated. Genotypes were coded on a scale of 0 to 4. Root mean squared error (RMSE) was calculated between actual genotypes and genotypes ascertained from simulated RAD-seq reads at 395 SNP markers (lower RMSE = higher accuracy). Each point represents one SNP. Median read depth is indicated by color, including genotypes with zero reads. The RMSE for continuous genotypes output by the polyRAD PopStruct LD method is shown on the x-axis, and the RMSE of other methods and types of genotypes (continuous or discrete) is shown on the y-axis. The dashed line indicates the ordinary least-squares regression with slope and intercept estimates, with standard errors. The “norm” model was used with updog. (A) RMSE calculated using only genotypes with more than zero reads. (B) RMSE calculated using only genotypes with zero reads, by genotyping or imputation method and genotype type. LinkImpute was not included given that it works for diploids only.

In diploid *M. sinensis* and tetraploid potato F1 mapping populations, polyRAD outperformed the GATK method, fitPoly, and updog, particularly when linked markers were used for informing the priors in polyRAD (Figs. 4A and 5A). In diploids and tetraploids respectively using genotypes with non-zero read depth, error (RMSE) using the polyRAD linkage model with discrete genotypes was reduced by 31.6% (SE 2.2%) and 48.0% (SE 0.4%) with respect to the GATK model, and 1.5% (SE 3.1%) and 17.1% (SE 0.6%) with respect to the updog “f1” model with discrete genotypes. For diploids, error was reduced by 39.8% (SE 2.5%) using polyRAD with respect to fitPoly, while for tetraploids fitPoly failed for all markers. For imputation, polyRAD using the linkage model performed similarly to LinkImpute and rrBLUP (Figs. 4B and 5B). Although only F1 populations are presented here, many other population types are supported in polyRAD.

**Fig. 4.**
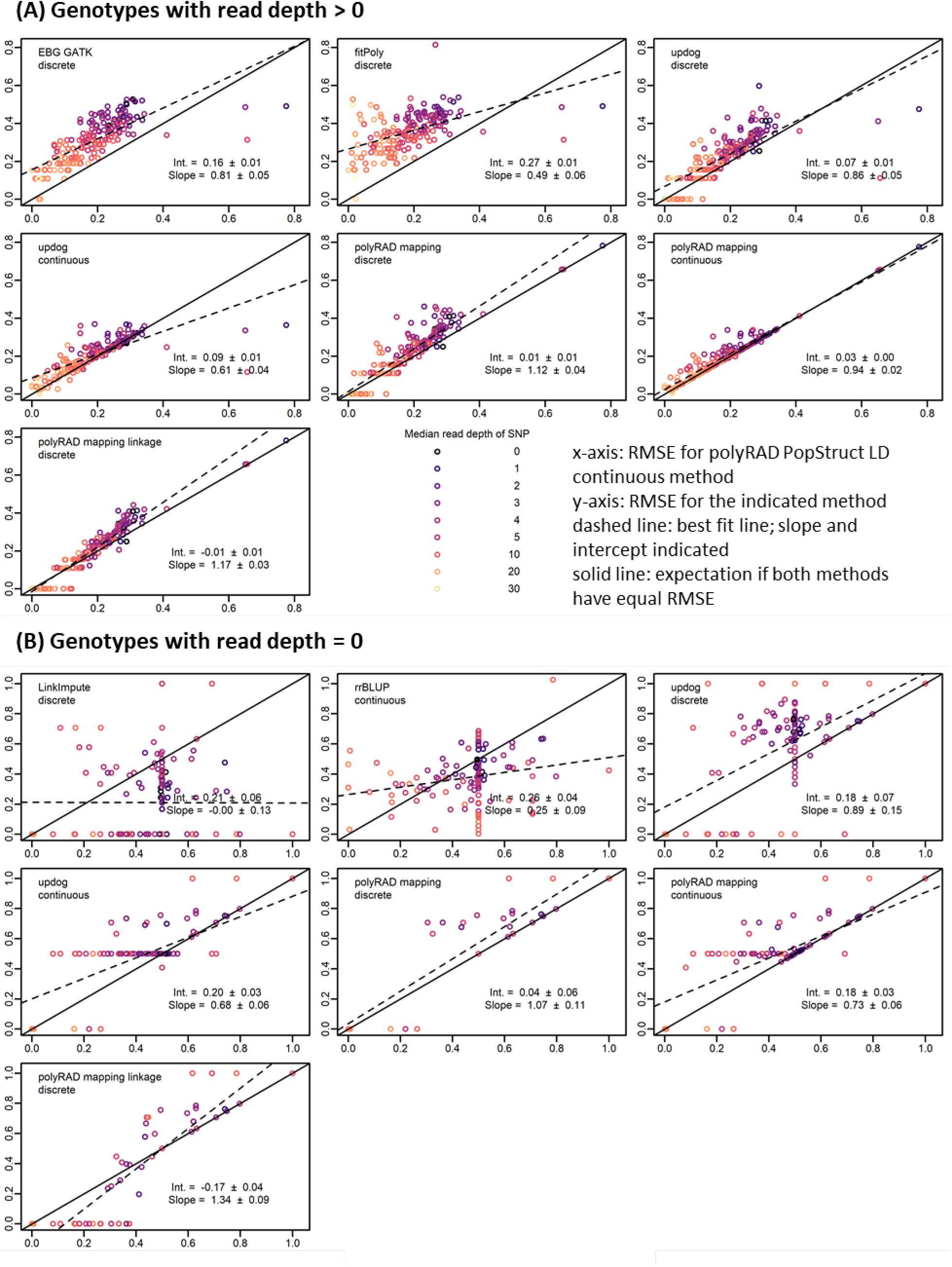
Genotyping error of EBG, fitPoly, updog, polyRAD, LinkImpute, and rrBLUP in an F1 mapping population of 83 diploid *Miscanthus sinensis*. The benefits of incorporating linkage into the genotyping model and using continuous rather than discrete genotypes are illustrated. Genotypes were coded on a scale of 0 to 2. Root mean squared error (RMSE) was calculated between actual genotypes and genotypes ascertained from simulated RAD-seq reads at 241 SNP markers (lower RMSE = higher accuracy). Each point represents one SNP. Median read depth is indicated by color, including genotypes with zero reads. The RMSE for continuous genotypes output by the polyRAD PopStruct LD method is shown on the x-axis, and the RMSE of other methods and types of genotypes (continuous or discrete) is shown on the y-axis. The dashed line indicates the ordinary least-squares regression with slope and intercept estimates, with standard errors. The “f1” model was used with updog. (A) RMSE calculated using only genotypes with more than zero reads. (B) RMSE calculated using only genotypes with zero reads, by genotyping or imputation method and genotype type.

**Fig. 5.**
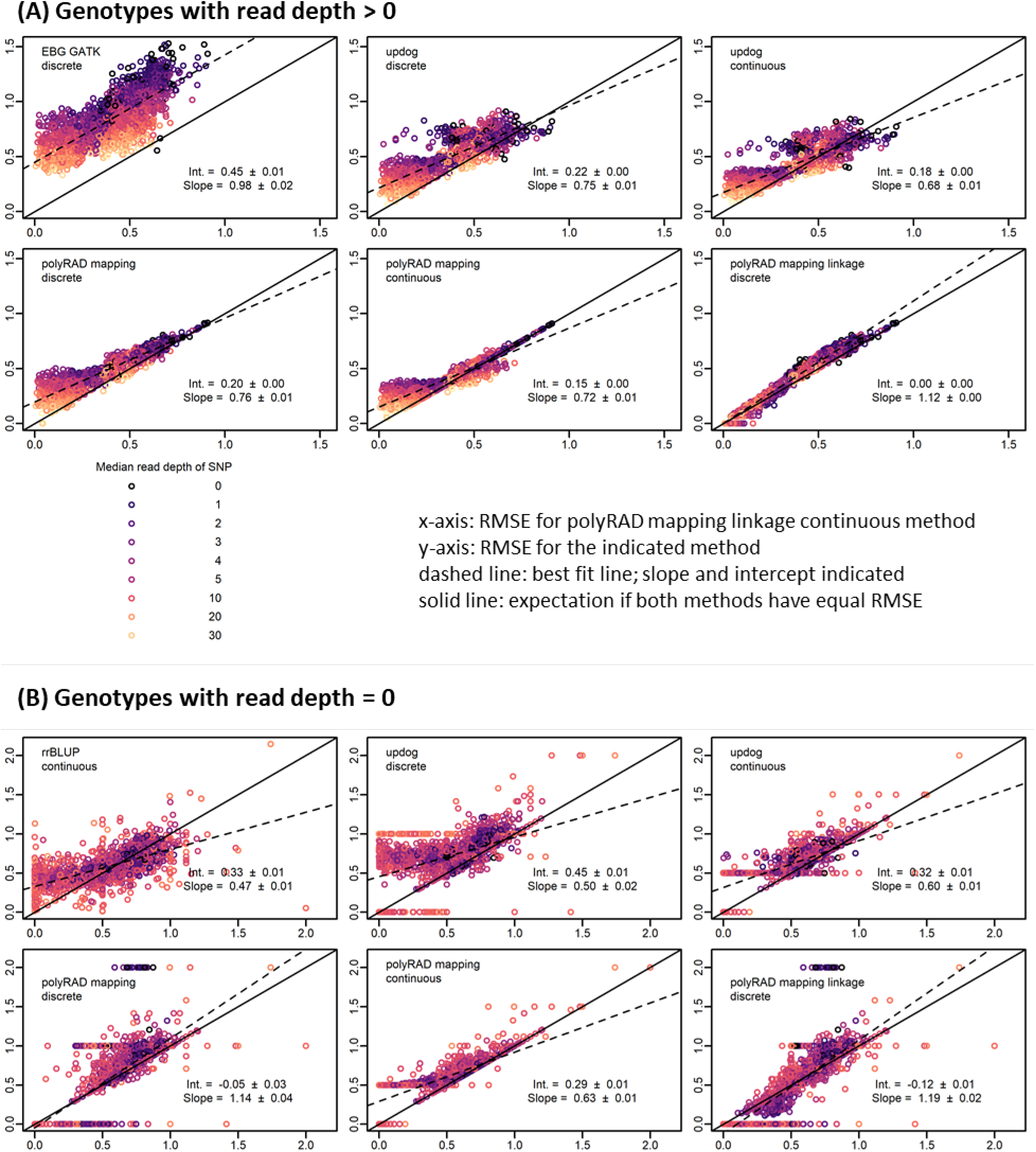
Genotyping error of EBG, updog, polyRAD, and rrBLUP in an F1 mapping population of tetraploid potato with 238 progeny. The benefits of incorporating linkage into the genotyping model and using continuous rather than discrete genotypes are illustrated. Genotypes were coded on a scale of 0 to 4. Root mean squared error (RMSE) was calculated between actual genotypes and genotypes ascertained from simulated RAD-seq reads at 2538 SNP markers (lower RMSE = higher accuracy). Each point represents one SNP. Median read depth is indicated by color, including genotypes with zero reads. The RMSE for continuous genotypes output by the polyRAD mapping method with linkage is shown on the x-axis, and the RMSE of other methods and types of genotypes (continuous or discrete) is shown on the y-axis. The dashed line indicates the ordinary least-squares regression with slope and intercept estimates, with standard errors. The “f1” model was used with updog. fitPoly results are omitted since it failed for all markers, and LinkImpute was not run since LinkImpute is for diploids only. (A) RMSE calculated using only genotypes with more than zero reads. (B) RMSE calculated using only genotypes with zero reads, by genotyping or imputation method and genotype type.

Genotyping error was also reduced 10-15% in most cases by exporting genotypes as continuous numerical variables (posterior mean genotypes), rather than discrete values (Figs. 2-5). For example, in a diploid, a true heterozygote (numeric value of 1) with reads only for the reference allele might erroneously be called as zero (homozygous for the reference allele) if only the most probable genotype is exported. However, the genotype could be called 0.4 if continuous genotypes are allowed, indicating that there is a 60% chance of it being a homozygote and 40% chance of it being a heterozygote, and thereby reducing the error from 1.0 to 0.6. Similarly in polyploids, continuous numerical genotypes can correct for errors in allele copy number estimation of heterozygotes.

### Downstream applications and implications for sequencing strategies

The genotyping methods implemented in polyRAD will have the most benefit for marker analysis where 1) the accuracy of individual genotypes is important, and 2) genotypes can be treated as continuous rather than discrete variables. The use of continuous versus discrete genotypes has been demonstrated to increase power for genome-wide association studies (GWAS) (Grandke *et al*. 2016) and genomic prediction (Oliveira *et al*. 2018) in polyploids. More generally, we anticipate that analyses that seek to quantify marker-trait associations in a population of individuals, including GWAS, quantitative trait locus mapping, and genomic prediction methods involving variable selection, will especially benefit from polyRAD. By reducing genotyping error, polyRAD will increase the power of these methods to detect true associations. Analyses that will benefit less from polyRAD genotyping are those where an average is taken across many genotypes in order to estimate a statistic, such as allele frequencies in a population or overall relatedness of individuals (including kinship-based methods of genomic prediction), because genotyping errors generally are not biased towards one allele or the other and tend to balance out over many individuals and loci (Buerkle and Gompert 2013; Dodds *et al*. 2015).

The advantages of polyRAD for accurate genotyping at low sequence read depth alter the economics of sequence-based genotyping, enabling researchers to purchase fewer sequencing lanes, multiplex more samples per lane, and/or retain more markers during filtering. In particular, for protocols using restriction enzymes where read depth varies considerably from locus to locus depending on fragment size (Beissinger *et al*. 2013; Davey *et al*. 2013; Andrews *et al*. 2016), there are diminishing returns on increasing the per-sample read depth, because some loci receive far more reads than are needed for accurate genotyping while other loci remain poor quality. Using population structure and linkage between loci, polyRAD uses information from high-depth markers to improve genotyping accuracy of low-depth markers, helping to maximize the useful information that is obtained from sequencing data. This advance is expected to improve breeding efficiency and economics.

## Acknowlegements

This material is based upon work supported by the National Science Foundation under Grant No. 1661490. The authors thank Marcus Olatoye, Per McCord, and Joyce Njuguna for testing the polyRAD software, and Associate Editor Daniel Runcie and two anonymous reviewers for comments on an earlier version of this manuscript.

